# Annotating Macromolecular Complexes in the Protein Data Bank: Improving the FAIRness of Structure Data

**DOI:** 10.1101/2023.05.15.540692

**Authors:** Sri Devan Appasamy, John Berrisford, Romana Gaborova, Sreenath Nair, Stephen Anyango, Sergei Grudinin, Mandar Deshpande, David Armstrong, Ivanna Pidruchna, Joseph I. J. Ellaway, Grisell Díaz Leines, Deepti Gupta, Deborah Harrus, Mihaly Varadi, Sameer Velankar

**Author notes:** contributed equally to this work (Sri Devan Appasamy, John Berrisford).

## Abstract

Macromolecular complexes are essential functional units in nearly all cellular processes, and their atomic-level understanding is critical for elucidating and modulating molecular mechanisms. The Protein Data Bank (PDB) serves as the global repository for experimentally determined structures of macromolecules. Structural data in the PDB offer valuable insights into the dynamics, conformation, and functional states of biological assemblies. However, the current annotation practices lack standardised naming conventions for assemblies in the PDB, complicating the identification of instances representing the same assembly.

In this study, we introduce a method leveraging resources external to PDB, such as the Complex Portal, UniProt and Gene Ontology, to describe assemblies and contextualise them within their biological settings accurately. Employing the proposed approach, we assigned standard names and provided value-added annotations to over 90% of unique assemblies in the PDB. This standardisation of assembly data enhances the PDB, facilitating a deeper understanding of these cellular components. Furthermore, the data standardisation improves the PDB’s FAIR attributes, fostering more effective basic and translational research and education across scientific disciplines.

## Background and Summary

Macromolecular complexes (or assemblies) composed of proteins and nucleic acids are integral to nearly all cellular processes. Assemblies such as RNA polymerase and ribosome are key players in the transmission of genetic information from DNA to proteins, by transcribing genetic information stored in DNA into RNA and translating RNA-encoded information into proteins, respectively (1, 2). Assemblies can be broadly classified into stable and transient complexes (3). Large assemblies, such as ribosomes, exhibit high stability, while others form transient yet stable assemblies depending on the cellular context, as seen in signalling pathways (4) (Figure 1). Determining the 3D structures of these assemblies is crucial for understanding their functional mechanisms and designing novel therapeutics targeting these molecular machines (5–7).

**Figure 1.**
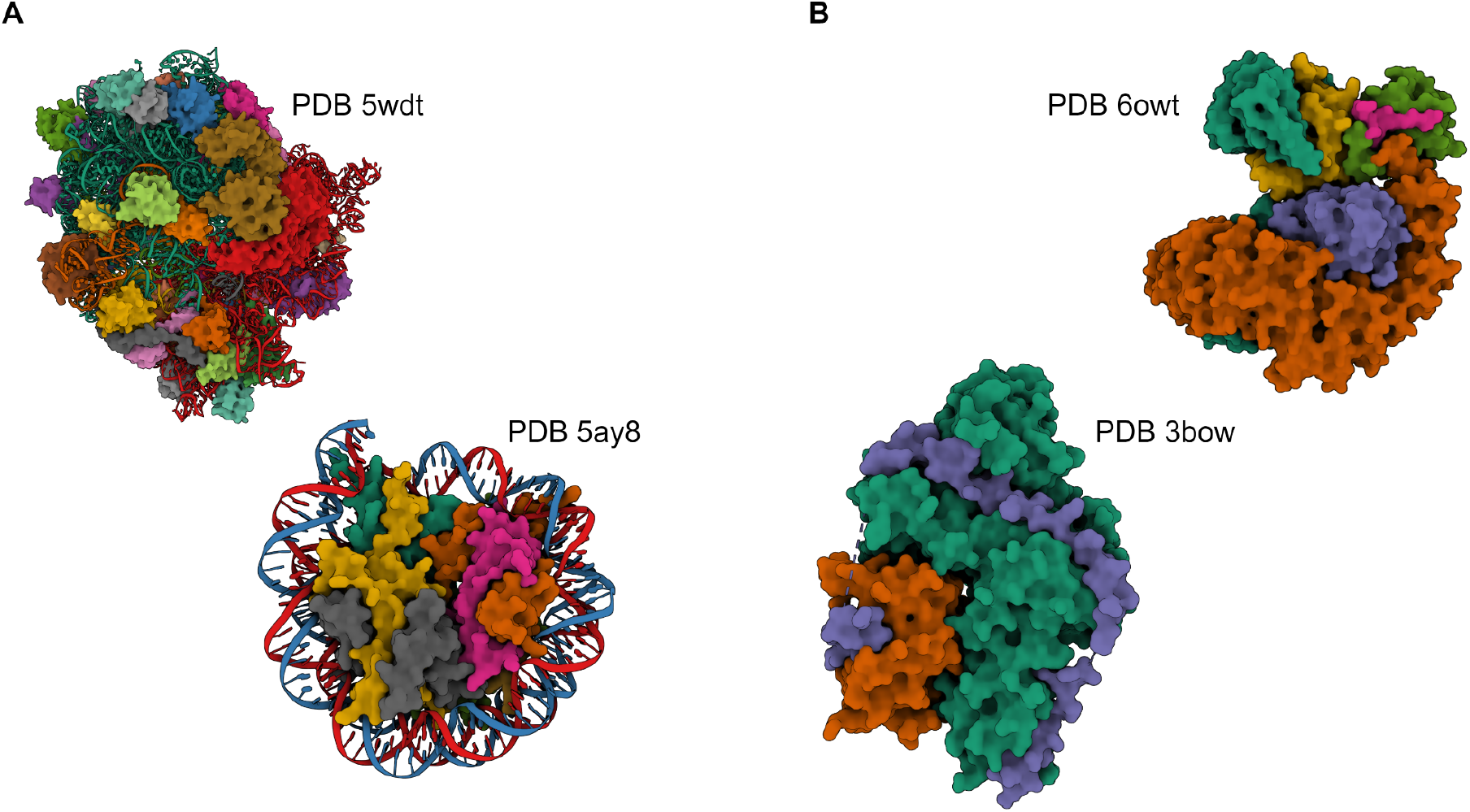
Stable and transient biological complexes. Bacterial ribosomes (PDB 5wdt) and the human nucleosome (PDB 5ay8) are examples of stable macromolecular machines (panel A). The clathrin adaptor AP-2 complex (PDB 6owt) and the calpain-calpastatin complex (PDB 3bow) are examples of transient complexes.

X-ray crystallography, cryogenic-electron microscopy (cryo-EM), and nuclear magnetic resonance spectroscopy (NMR) are essential techniques for experimentally determining macromolecular assembly structures. X-ray crystallography has elucidated structures of various assemblies, including ATP synthase (8–10), membrane proteins (11–13), proteasomes (14–16), and ribosomes (17–19). NMR has provided insights into structures such as the Hsp90-Tau complex (20), the box C/D ribonucleoprotein enzyme complex (21) and several molecular chaperones (22, 23). Recent advances in structural biology, particularly cryo-EM (24, 25) and integrative hybrid methods (26) have enabled routine investigation of larger macromolecular assemblies like nuclear pore complex (27), BBsome complex (28) and mammalian circadian clock complexes (29).

The Protein Data Bank (PDB) (30) is the unified, global repository for experimentally determined macromolecular structures. The worldwide PDB (wwPDB) organisation manages the archive, ensuring free and public access to structural data for the global community (31). The Electron Microscopy Data Bank (EMDB; 32) is a public repository for cryo-microscopy electric potential maps and tomograms of macromolecular complexes and subcellular structures. The Small Angle Biological Data Bank (SASBDB; 33) is a curated repository for bio-macromolecular small-angle scattering of X-rays and neutrons (SAXS and SANS) data and models. The Biological Magnetic Resonance Bank (BMRB; 34) archives spectral and quantitative data from NMR spectroscopic investigations of biological macromolecules. Lastly, PDB-DEV (35) is a prototype archiving system for structural models obtained using integrative or hybrid modelling approaches.

The PDB archive may house multiple structures representing the same assembly, offering unique opportunities to study structure ensembles and gain a mechanistic understanding of large macromolecular assemblies. Molecular structures can be determined under various conditions to explore the conformational space or obtain functional insights. Researchers may introduce engineered mutations, modify pH and ion concentrations, or add molecules ranging from ligands to antibodies. Additionally, structures may be solved at different resolutions or in distinct space groups. Multiple assemblies may be derived from the same crystal form describing various potential quaternary structures. For instance, PDB entry 1e94 (HslV-HslU protease complex) features one assembly representing a homo-hexamer and another a homo-dodecamer.

To address this ambiguity, the Protein Data Bank in Europe (PDBe) (36) defines the preferred assembly as the smallest assembly containing all polymeric entities. This approach enables users to identify the most probable assembly observed in the experiment represented in the PDB entry for a given macromolecule, which is also displayed in PDBe’s search results. In the HslV-HslU complex example, the homo-hexamer form is displayed as the preferred assembly.

Identifying and analysing protein assemblies involves distinguishing biologically relevant interfaces from protein-protein contacts caused by crystal packing (37–39). Several methods have been developed to address this issue, such as PISA (Protein Interfaces, Surfaces, and Assemblies) (37), which employs thermodynamic estimation of interface stability, and EPPIC (Evolutionary Protein-Protein Interface Classifier) (40), which uses evolutionary information from protein sequences to differentiate biological interfaces from lattice contacts. Another method, QSalign (41), identifies biologically relevant assemblies by structurally aligning quaternary structures and inferring ones with conserved interfaces as biologically relevant. These three methods were integrated into a single predictor called QSbio, which provides a confidence score for biological relevance of given assemblies (41).

Finding instances representing the same assembly in the PDB is challenging due to current annotation practices, which do not include consistent naming and analysis of observed complexes across the PDB archive. For instance, the assembly in PDB entry 6kat is not described as haemoglobin but as a complex of two haemoglobin subunit alpha chains and two haemoglobin subunit beta chains. Additionally, since a given set of coordinates and space group symmetry described in a PDB entry can result in multiple assemblies with different stoichiometry or components, it is not easy to identify the correct biologically relevant assembly and the entry title does not necessarily reflect these directly. Therefore, the lack of consistent naming makes it difficult to find whole or partial complexes through PDB searches.

To address this issue, we have compiled all unique assemblies in the PDB by identifying individual components using their mappings to external resources, such as UniProt for proteins through the SIFTS resource (42) and Rfam (43) for RNA molecules. We then determined the stoichiometry of individual components within a complex. Based on this process, we generated a list of PDB entries corresponding to each assembly with unique composition. Establishing this mechanism and assigning stable identifiers for each unique assembly across the PDB archive will promote the study and understanding of conformational changes and molecular mechanisms. In addition to stable identifiers, we have assigned human-readable, and in some cases, manually curated names and mapped the PDB assemblies to Complex Portal (44) entries where possible. The assembly identification and the naming process are incorporated into PDBe’s weekly release cycle, ensuring data integrity, persistence and concurrency.

## Results

### Composition of assemblies in the PDB

The identification of unique assemblies in the PDB is based on the preferred assemblies defined using the process described in the Methods section below. These encompass monomeric proteins (e.g., lysozyme) and higher-order preferred assemblies of homomeric or heteromeric complexes (e.g., viruses or ribosomes). Of the 97,528 unique assembly compositions in the PDB excluding those containing chimeric chains (as of late March 2023), 81,792 are protein-only, 11,613 are protein-nucleic acid assemblies, and the remaining 4,123 consist of nucleic acid only (Figure 2). Henceforth, the term “unique assemblies” refers to the set of unique PDB assemblies based on the composition of each assembly. Additionally, 38,789 of these assemblies are heteromeric assemblies, 31,526 are homomeric assemblies and the rest are monomeric assemblies. In 74,427 of the unique assemblies, at least one component can be mapped to a UniProt accession, and in 63,894 assemblies, all polymer components map to UniProt accessions.

**Figure 2.**
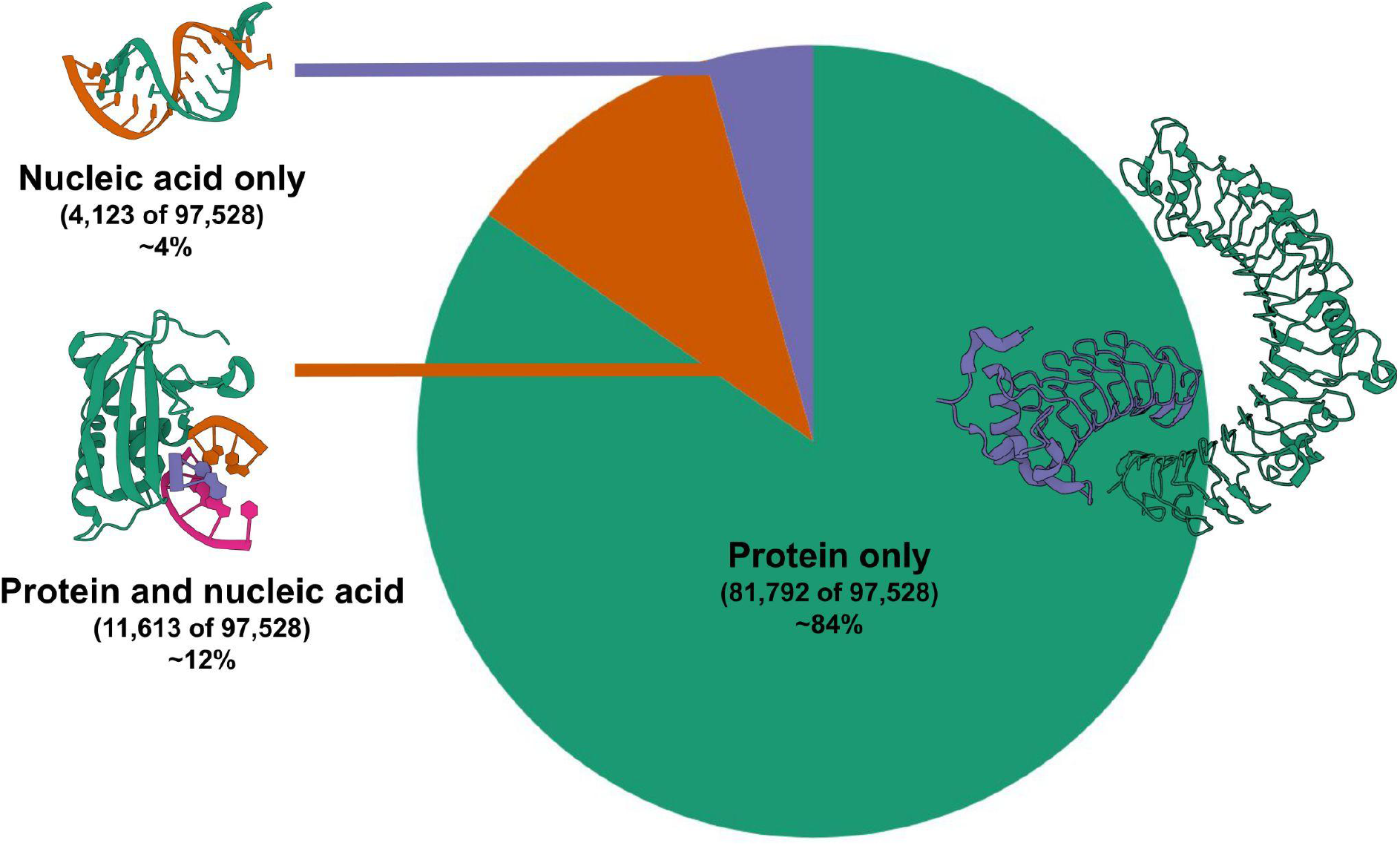
Assembly composition in the PDB. Protein-only assemblies dominate the macromolecular assemblies, and most proteins can be mapped to UniProt accessions. Example PDB entries from protein-only, protein-nucleic acid and nucleic acid-only assemblies include PDB entries 6bxa, 6dpo and 6c8m, respectively.

X-ray diffraction is the most common method used to determine the structures of unique assemblies (77,984), followed by electron microscopy (9,369) and nuclear magnetic resonance (7,902), respectively (Table 1). A total of 1,556 unique assemblies have structures solved using both NMR and X-ray diffraction, 470 using both EM and X-ray diffraction, and 41 were solved using all three different methods (Figure 3). Additionally, we identified the species for each unique assembly whenever possible. The top ten species include bacteria and eukaryotic organisms, with human assemblies dominating the list (Table 2).

**Table 1.** Top five experimental methods or combinations of experimental methods used to solve structures of unique assemblies. X-ray diffraction is the predominant method for determining the structures of assemblies, followed by electron microscopy (EM) and nuclear magnetic resonance (NMR).

| Experimental Method | Unique Assemblies |
| --- | --- |
| X-ray diffraction | 77,984 |
| Electron microscopy | 9,369 |
| Nuclear magnetic resonance | 7,902 |
| Nuclear magnetic resonance, X-ray diffraction | 1,556 |
| Electron microscopy, X-ray diffraction | 470 |

**Table 2.** Top ten species from which unique assemblies are solved. Number of unique assemblies in the PDB containing human proteins, followed by bacteria and model organisms.

| Scientific Name | Unique Assemblies |
| --- | --- |
| *Homo sapiens* | 16,645 |
| *Escherichia coli* | 3,721 |
| *Mus musculus* | 2,940 |
| *Saccharomyces cerevisiae* | 2,778 |
| *Thermus thermophilus* | 1,154 |
| *Pseudomonas aeruginosa* | 1,086 |
| *Mycobacterium tuberculosis* | 1,050 |
| *Arabidopsis thaliana* | 995 |
| *Mus norvegicus* | 989 |
| *Mycobacterium tuberculosis* | 900 |

**Figure 3.**
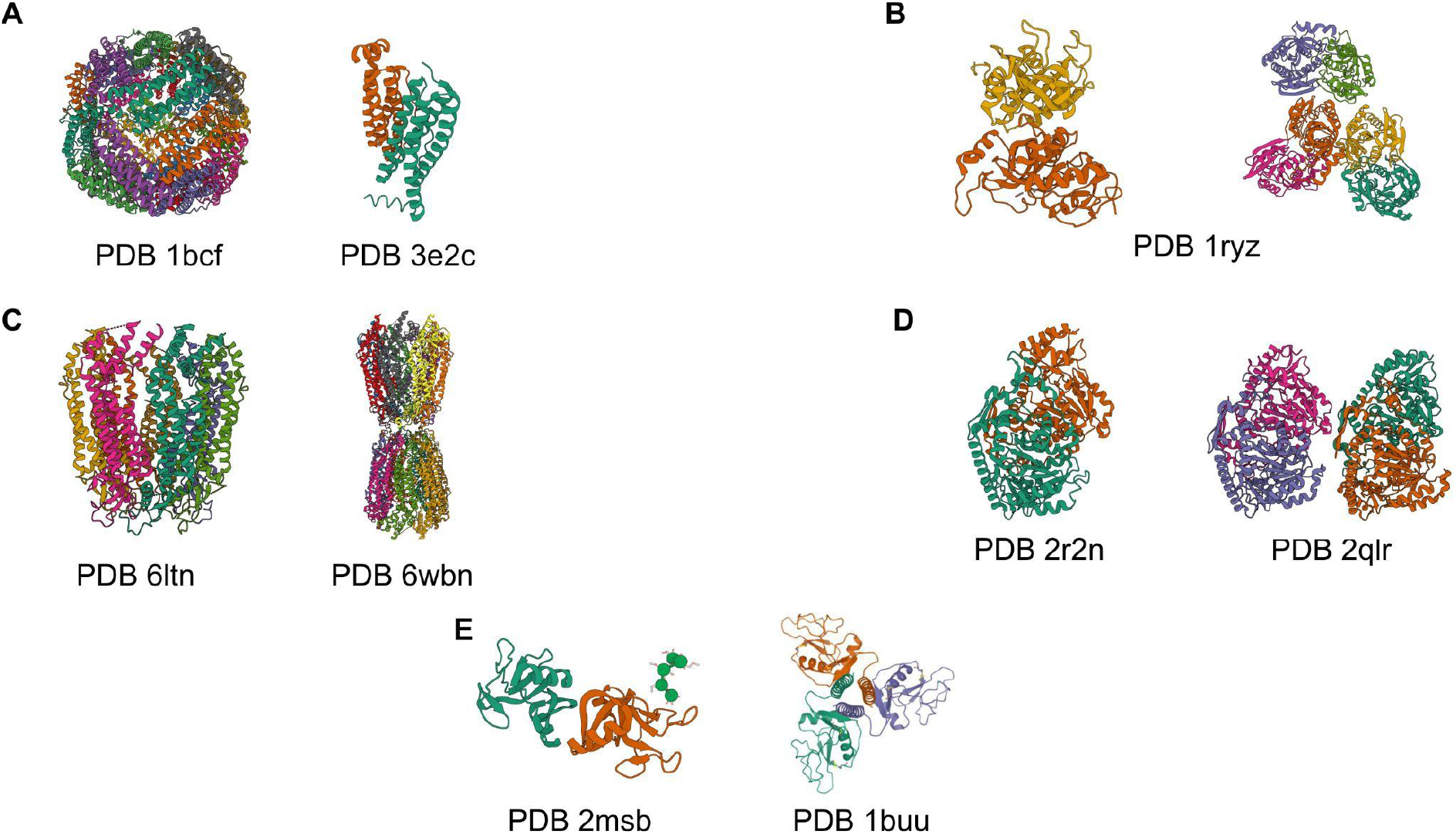
Examples of homomeric assemblies with different stoichiometries in the PDB. We identified five main reasons for observing multiple stoichiometries for an assembly. These differences can be caused by experimental conditions (panel A), difficulties in automated assembly assignments (panel B), challenges in the curation and annotation process of assemblies (panel C), genuine errors in curation (panel D), and differences in the sample, for example in the sequence length (panel E).

To uniquely identify assemblies, we combined mappings to external resources with the stoichiometry of the components. Assemblies with components that map to the same UniProt accessions but have different stoichiometries may indicate varying experimental conditions or incorrect annotation. Analysis of nearly 28,000 homomeric protein assemblies, where all components can be mapped to UniProt, revealed 2,955 cases with different stoichiometries. Table 3 displays some of the proteins where the homomeric assemblies exhibit highly variable compositions.

**Table 3.** Homomeric proteins with multiple stoichiometries in the PDB.

| UniProt accession | Protein name | Number of different stoichiometry |
| --- | --- | --- |
| P05067 | Amyloid-beta precursor protein | 12 |
| P68135 | Actin, alpha skeletal muscle | 10 |
| P10636 | Microtubule-associated protein tau | 10 |
| P10997 | Islet amyloid peptide | 9 |
| P37840 | Alpha-synuclein | 8 |
| B2J6D9 | Phage shock protein A, PspA | 7 |
| P02766 | Transthyretin | 7 |
| P04156 | Major prion protein | 7 |
| P41784 | Protein PrgI | 7 |
| Q13148 | TAR DNA-binding protein 43 | 7 |

The PDB is an extensive resource of protein structures, containing over 200,000 entries. However, this abundance of information presents challenges in curating and annotating the data. Proteins can exist in various stoichiometries, such as dimers, tetramers, and larger oligomers, and thus, selecting the correct assembly state or more commonly known as the biological assembly in a crystal structure can be challenging as experimental conditions and protein constructs may alter the oligomeric state of protein during structure determination. Furthermore, errors in the curation process may lead to incorrect assembly states being assigned. Our analysis of homomeric protein assemblies highlights several examples of proteins with different stoichiometries in the PDB, the challenges associated with determining the correct assembly state, and some instances of inaccuracies (Figure 3).

An example of a protein with multiple stoichiometries in the PDB, explainable by experimental conditions, is bacterioferritin (UniProt accession P0ABD3), which typically forms a 24-meric sphere (e.g., PDB 1bcf) but in some PDB entries, an engineered version forms dimers (e.g., PDB 3e2c). However, other cases exemplify the challenges of automatically selecting the representative assembly for a given PDB, such as PDB entry 1ryz, which includes four curated assemblies. The first two represent different hexameric forms, while the latter represents alternative dimeric forms. The PDBe process for selecting preferred assemblies favours one of the dimeric forms, but the QSbio assembly predictor assigns the highest confidence to the hexamer.

The curation and annotation of PDB entries present significant challenges, as illustrated by several examples. For instance, the protein Histone-arginine methyltransferase CARM1 (UniProt accession Q9WVG6) is represented as a homo-tetramer in 30 PDB entries (e.g. PDB entry 5ih3) and as a homodimer in 2 PDB entries (e.g. PDB 2v74). The homodimers in PDB entry 2v74 have a 98% confidence level, according to the QSbio predictor. Similarly, human Pannexin1 (UniProt accession Q96RD7) is a homo-heptamer in 18 PDB entries (e.g. PDB entry 6ltn) and a homo-tetradecamer in one example (PDB entry 6wbn). The publication associated with the homo-tetradecamer structure suggests that this assembly arrangement does not result from incorrect curation (45).

Differences in sample sequence when solving the assembly structure can also result in multiple stoichiometries for an assembly. For example, the mannose-binding protein forms a trimer in its active form (e.g. PDB 1buu). However, PDB entry 2msb represents a deviation from the usual stoichiometry as it forms a dimer due to its shorter sample sequence that does not contain the necessary helix that forms the trimer interface.

Our analysis revealed a few incorrect assembly annotations, such as human Kynurenine/alpha-aminoadipate aminotransferase (UniProt accession Q8N5Z0). This protein is a homodimer in 15 PDB entries (e.g. PDB 2r2n) and a homotetramer in only one PDB entry (PDB 2qlr). The QSbio predictor suggests a 98% confidence level for the homodimer, highlighting the need for careful curation and annotation.

### Sub- and super-assemblies

Biological assemblies can have multiple components, each with varying stoichiometry. The ribosome, responsible for protein synthesis in every living cell, exemplifies this variability. The ribosome comprises two ribosomal subunits and can bind multiple tRNAs, mRNA, and diverse protein factors. The binding of tRNAs and various protein factors can induce large-scale conformational changes in the ribosome, affecting its function (46, 47). The empty ribosome (e.g., PDB 4ybb) can be considered a subassembly compared to the ribosome with bound tRNAs or protein factors (e.g., PDB 5uym). Identifying sub- and super-assemblies is key to the identification of the transient assemblies and understanding each component’s contribution to the function of the assembly and the relationships between different components.

Our proposed approach allows us to identify sub- and super-assemblies in the PDB. We found that over 5% of assemblies in the PDB are sub-assemblies (5,099 out of 97,528). Over 40% of these sub-assemblies are components of two or more assemblies (2,118 out of 5,099). For example, lysozyme (e.g., PDB 6kd1), one of the most common proteins in the PDB, has 24 super-assemblies containing additional components that map to UniProt, such as fibronectin (e.g., PDB 5j7c), aspartate-tRNA ligase (e.g., PDB 4gla), and periplasmic pH-dependent serine endoprotease DegQ (e.g., PDB 4a8a). Similarly, around 15% of the assemblies in the PDB are super-assemblies of another assembly (14,279 out of 97,528).

### Human-readable names for assemblies in the PDB

The Complex Portal (44) provides recommended names for macromolecular complexes with defined components and stoichiometry from selected model organisms. Currently, the Complex Portal contains around 4,000 annotated complexes, representing over 5% of the unique complexes in the PDB (70,315). When comparing the composition of assemblies in the PDB to those in the Complex Portal, we found that 1,648 unique PDB complexes match the exact compositions in Complex Portal. Extending the mapping to include PDB assemblies with additional components increases the number of matches between PDB and Complex Portal to 2,129 PDB assemblies. Of these, 371 contain an additional protein component, 67 have an additional DNA component, and 13 include an additional RNA component.

For example, the PDB contains three assemblies of the Cyclin A2-CK2 complex with Cyclin-dependent kinase inhibitor 1B (PDB 1h27, 1jsu, and 6ath). Since the core Cyclin A2-CK2 complex maps to Complex Portal ID CPX-2006, these assemblies can be labelled by combining the name in Complex Portal with an additional protein component (E.g. Cyclin A2-CK2 complex and Cyclin-dependent kinase inhibitor 1B). A further 344 assemblies contain components that match a Complex Portal definition and have additional components mapped to other Complex Portal entries. An example is PDB entry 6r8z, which contains Nucleosome variant H3.1-H2A.2-H2B.1, UV DNA damage recognition assembly DBB1-DBB2 with DNA and maps to two Complex Portal accessions CPX-2556 and CPX-308, respectively. Following these conventions allowed us to name 2,473 unique PDB assemblies using names obtained from Complex Portal. We have made the generated mapping files available to the Complex Portal team and the broader scientific community through the public PDBe-KB FTP area, which is available at https://ftp.ebi.ac.uk/pub/databases/pdbe-kb/complexes/.

For protein assemblies that comprise a single component, we can use the name of that component regardless of stoichiometry. Since almost two-thirds of protein assemblies in the PDB are either monomeric or homomeric, we could use the protein name from UniProt to name these assemblies. Using this approach, we could name nearly 70% of the assemblies in the PDB. The remaining assemblies contain additional components, such as nucleic acids, antibodies, or peptides. In these cases, we can assign names by combining the name of the protein with a generic component label. For example, the assembly in PDB entry 5hi4 can be named “Interleukin-17A, IG-heavy chain, IG-light lambda chain, peptide complex” by combining the Interleukin-17A name from UniProt with the antibody and peptide names. In addition, we used a combination of antibody and molecule names to name another 7,097 protein assemblies that consist of unmapped components.

We have also used the Gene Ontology (GO) cellular component terms to name complexes since they describe the whole complex in cases where the individual components have GO annotations. Using this approach, we can name an assembly if every component of the PDB assembly is annotated with a common GO term. Applying this approach to the PDB archive, we named 400 unique assemblies. These assemblies include 132 proteasome complexes, 32 photosystem II complexes, 27 photosystem I complexes, 14 haemoglobin complexes, and 8 ATP synthase V complexes

We also identified and named another 259 unique PDB assemblies representing biological complexes using a manually curated list prepared by PDBe curators. By combining the naming approaches described above, we could name over 90% (90,999 out of 97,528) of assemblies in the PDB (Table 4).

**Table 4.** Breakdown of names assigned to unique assemblies based on our naming approach.

| Naming category | Number of unique assemblies | Number of annotated PDB entries |
| --- | --- | --- |
| UniProt | 66,110 | 166,096 |
| Heterodimer | 8,206 | 13, 749 |
| Unmapped protein (excluding antibodies) | 5,542 | 5,542 |
| General nucleic acid name (DNA, RNA or DNA/RNA) | 3,893 | 3,893 |
| Complex Portal | 2,457 | 6,746 |
| Ribosome | 1,582 | 1,608 |
| Antibody | 1,555 | 1,555 |
| Common name from entity names | 893 | 2,070 |
| Gene Ontology | 400 | 858 |
| PDBe curated | 259 | 616 |
| Rfam (excluding ribosome) | 102 | 336 |

The bulk of the remaining unnamed assemblies (∼6,000) are largely heteromeric assemblies containing both mapped and unmapped components that cannot be named automatically using our current process. We are working to create new rules and using manual curation for naming these assemblies.

### Finding ribosomes in the PDB

Complexes in the PDB are often heterogeneous in the composition of subunits, with some instances missing components or having components that do not map to UniProt. This heterogeneity is most prevalent in large complexes, such as ribosomes, with various assemblies bound to diverse molecules.

We developed a combined approach to address this heterogeneity and enable the identification of ribosomes even when they lack certain ribosomal proteins or rRNA subunits. We identified ribosomes that contained both ribosomal RNAs mapped to Rfam and ribosomal proteins. In cases where a potential ribosome did not have mappings to Rfam, it was required to have both RNA and ribosomal protein components. By using this approach, we were able to identify a total of 1,582 unique ribosome assemblies.

To name each ribosome complex, we used either the name of the ribosomal subunit, such as 30S ribosomal subunit (e.g. PDB 6v3e), or the full ribosome name if both ribosomal subunits were present, such as 70S ribosome (e.g. PDB 5j7l). If the ribosome contained tRNA, identified through matching Rfam accessions RF00005 or RF01852, we added that information to the name. Any additional RNA molecules that are bound to the ribosome are simply named as RNA including messenger RNAs (mRNAs). For example, the PDB entry 6bok is an *E. coli* ribosome with both tRNA and mRNA bound and thus, is named as 70S ribosome and tRNA and RNA.

### Analysing the symmetry of assemblies

To infer the point-group symmetry operators of the members of each unique protein assembly composition based on the preferred assemblies, we used the AnAnaS software (48). This tool can detect five symmetry groups: cyclic, dihedral, tetrahedral, octahedral, and icosahedral. Among the symmetrical protein assemblies with unique compositions, cyclic (Cn, n≥2) and dihedral (Dn, n≥2) symmetry groups were the most frequent in the PDB, occurring in 77% and 20% of cases, respectively. Most cyclic symmetries were either C2 or C3, while the most common dihedral symmetries were D2, D3, and D4, in descending order of frequency (Figure 4).

**Figure 4.**
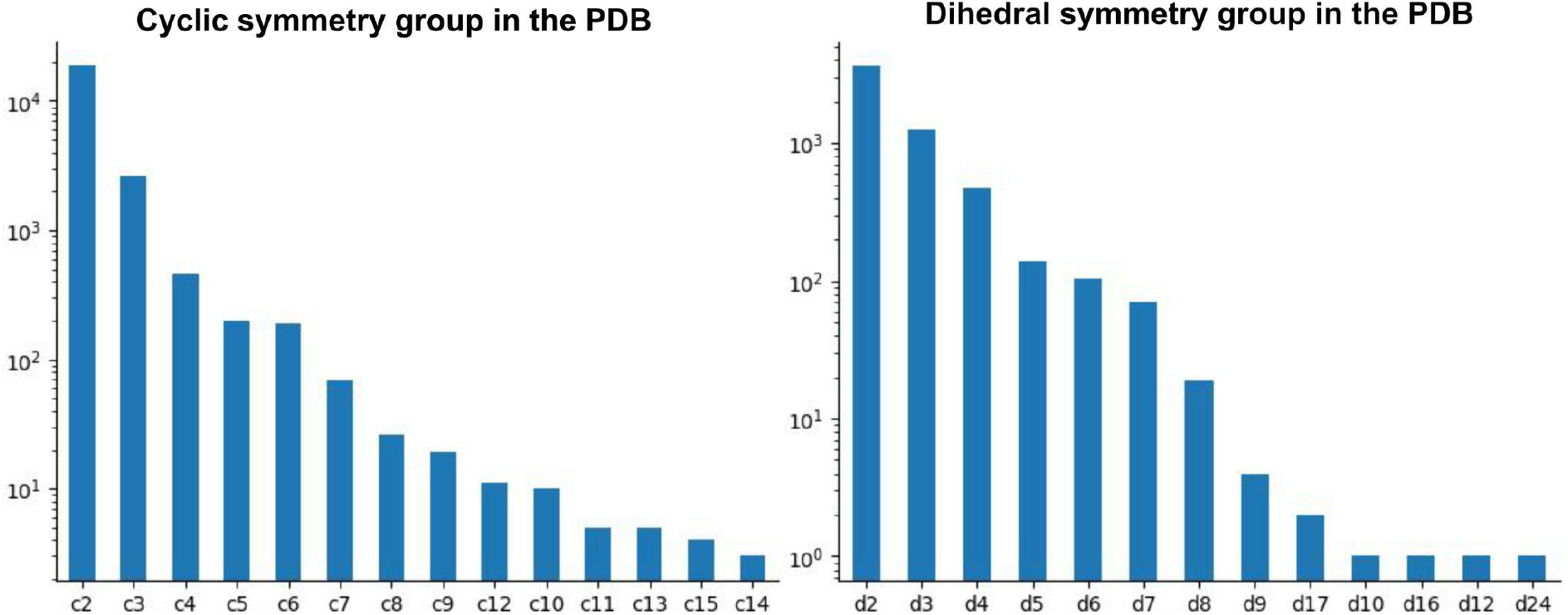
Frequency of cyclic and dihedral symmetries in the PDB. The PDB archive is dominated by cyclic c2 symmetry, with the second most frequent being dihedral d2 symmetry and cyclic c3 symmetry. The vertical axes in both plots are shown in logarithmic scale.

Our analysis revealed that close to 6.0% (1,651 out of 28,374) of these assemblies appear to have inconsistent symmetry operators among its members. For example, the PDB entry 4u7n is an inactive histidine kinase dimer (consists of two copies of UniProt A0A0M3KKX3) that has C2 symmetry. But other instances of this assembly (e.g. PDB entries 4u7o and 4zki) appear to lack symmetry. Individual domains of histidine kinases have been shown to adopt both symmetric and asymmetric conformations in different catalytic states which might explain the differences in the symmetry groups of these structures (49).

Symmetry analysis can also help detect unlikely assembly arrangements. For instance, the cytochrome P450 3A4 complex is formed as a homodimer in 35 PDB entries, with 29 PDB entries exhibiting C2 symmetry (e.g. PDB entry 5g5j). However, in PDB entries 7kvh, 7kvn, 7kvo, 7kvq, and 7kvs, there is no symmetry, and the arrangement of the monomers is different compared to other instances of the assembly due to very different crystal packing leading to a lack of symmetry. Another example is the homo-tetramer of L-asparaginase 2 (UniProt P00805) in PDB entry 6pa3, the only example among 45 PDB entries that does not have D2 symmetry in the provided biological assembly although the symmetry can be found in the crystal indicating a possible mistake in the annotation. In a few examples, symmetry is observed in one PDB entry, but most other entries do not have any symmetry. For example, RNA-binding protein Hfq (UniProt P0A6X3) is asymmetrical in PDB entry 4jrk but not in the 14 other examples of the same protein that exhibit c6 symmetry again indicating a probable issue with the identification of assembly that needs further investigation.

Based on our analysis, the AnAnaS software has proven useful in inferring symmetry operators for unique protein assembly compositions, with cyclic and dihedral symmetry groups being the most common in the PDB archive. A small percentage of these assemblies appear to have inconsistent symmetry operators requiring further investigation. However, analysing symmetry can help in the identification of unlikely assembly arrangements which may be biologically relevant.

## Discussion

Applying common data standards is crucial for making data Findable, Accessible, Interoperable, and Reusable (FAIR) (50). In this study, we have developed an automated process to identify over 90,000 unique assemblies in the Protein Data Bank by mapping individual components to external databases such as UniProt and Rfam. With the help of additional resources such as Complex Portal and Gene Ontology, we have assigned human-readable, consistent names to over 90% of these unique assemblies, addressing a significant deficiency in the curation process of structure data.

Our approach improves accessibility of structure data for a given complex via PDBe search mechanism. Previously, this type of search relied on the PDB entry title or the individual components in the assembly having a common naming convention, e.g. both Hemoglobin assembly components have Hemoglobin in their name, which makes it challenging to recover all the relevant information. The standardisation of assembly names and unique identifiers allow users to specify the PDBe complex identifier or the complex name under the advanced search option to find all the relevant information for a complex of interest. As shown in Figure 5, querying for Hemoglobin yields multiple variants of hemoglobin such as mutant adult human hemoglobin (PDB-CPX-159518), hemoglobin from parasitic flatworm *Fasciola hepatica* (PDB-CPX-163279) and foetal hemoglobin (PDB-CPX-159679). By clicking on the individual complex name, users can discover the same assembly across different species. Alternatively, clicking on the PDBe complex identifier would enable users to easily find all PDB entries containing a given assembly with identical composition and species.

**Figure 5.**
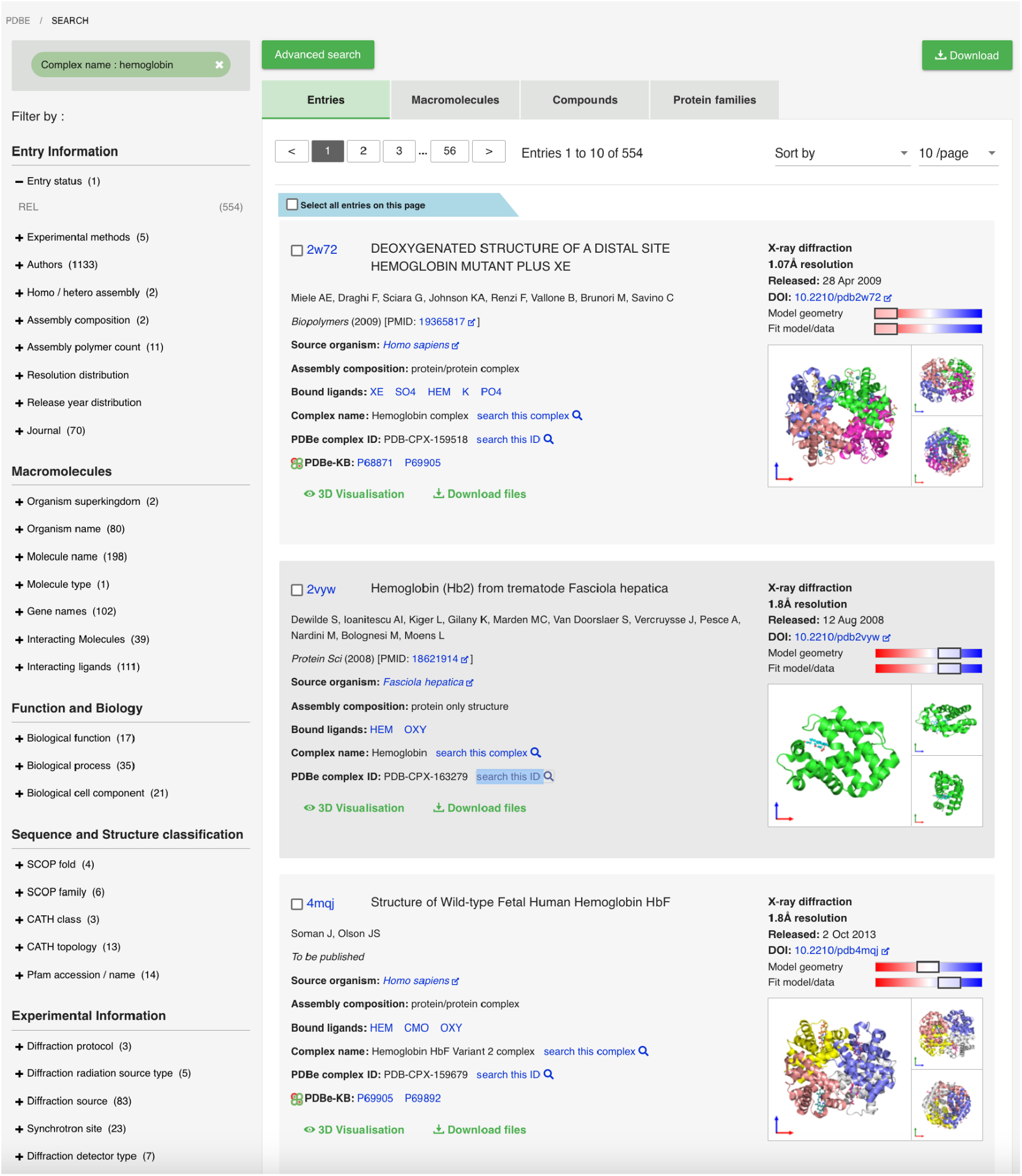
Finding complexes of interest in PDBe. By integrating the unique complex identifiers and complex names into the search system of PDBe, researchers can find distinct complexes more consistently across the PDB instead of relying on searching by PDB entry titles or complex component names.

Importantly, the consistent naming of assemblies, the assignment of stable accessions to each unique assembly, and integration with external reference annotations from UniProt, Rfam, GO, and Complex Portal have aided in putting these molecular machines in their biological contexts, facilitating the identification of meaningful relationships between function, assembly composition, and conformational heterogeneity. Furthermore, this high-quality, aggregated assembly data can serve as a training dataset and facilitate efforts to investigate the dynamics of macromolecular complexes computationally.

We believe that the identification and biocuration of all unique assembly compositions in the PDB archive have addressed a long-standing limitation of the archive and greatly enhanced the findability and reusability of these data to gain mechanistic insights into the biological context and function of these molecular machines. With consistently identified, unique assemblies labelled with human-readable names and stable accessions, users can easily search for specific assemblies and access all the relevant structure data for their analysis, advancing basic and translational research in life sciences. The implementation of our approach may serve as a model for improving the FAIRness of structural biology data.

## Methods

### Identification of unique assemblies

To create distinct assembly identifiers, we begin by attempting to map each component in the assembly to an accession/identifier from a reference database (refer to Table 5). UniProt accessions are used to map protein components, while mappings from the Rfam database (43) are used for RNA molecules. In addition, we also use antibody annotation generated by the tool ANARCI (51) to identify and name antibodies in the PDB. These accessions/identifiers are combined with stoichiometry numbers to generate description labels. We append the unmapped component type, PDB entry-id, entity-id, and stoichiometry to describe components that cannot be mapped to an external reference. We continue to work on classifying unmapped components and identifying suitable external references.

**Table 5.** Generating composition labels for assemblies in the PDB.

| Component type | Description format | Example |
| --- | --- | --- |
| Protein (UniProt) | [accession]_[stoichiometry] | P68871_1 |
| RNA (Rfam) | [accession]_[stoichiometry] | RF00177_1 |
| Antibody | antibody_[PDB]_[entity]_[stoichiometry] | antibody_5mv4_1_1 |
| Protein (unmapped) | protein_[PDB]_[entity]_[stoichiometry] | protein_7rx0_2_1 |
| RNA (unmapped) | RNA_[PDB]_[entity]_[stoichiometry] | RNA_1un6_2_1 |
| DNA | DNA_[PDB]_[entity]_[stoichiometry] | DNA_8b1t_4_1 |
| DNA/RNA hybrid | DNA/RNA_[PDB]_[entity]_[stoichiometry] | DNA/RNA_8e8j_3_1 |

To create a unique identifier for each complex, we combine the descriptions of the individual components and assign a distinct ID to each unique complex description. For instance, the description “P12345_1, DNA_1uty_2_1” is mapped to the identifier PDB-CPX-100487. We then compare the composition and stoichiometry of each assembly to those of complexes in the Complex Portal (44) and assign the relevant Complex Portal ID to the assembly whenever possible. To ensure the persistence of these IDs, we generate and store an md5 hash value for each unique complex composition, which serves as a reference for creating and maintaining persistent identifiers for all unique complexes in the PDB archive. At present, our approach only identifies assemblies within a given species and does not compare them across different species.

The next step in the process is to define sub-assemblies and super-assemblies based on the composition and stoichiometry data. We define a sub-assembly as an assembly where all its components are a subset of the components in another larger assembly. For instance, haemoglobin (two copies of UniProt P68871 and two copies of UniProt P69905) represented by the PDB entry 7jy3 is a subassembly of the haemoglobin-iron surface determinant B receptor complex (contains additional two copies of UniProt Q8NX66) represented by the PDB entry 7pch. We define a super-assembly as an assembly containing all the components of another assembly and additional members. Thus, in the previous example, the PDB entry 7pch is a super-assembly of haemoglobin (PDB entry 7jy3).

### Naming unique assemblies

To automate the naming of unique complexes, we developed a decision tree (Figure 6). First, we check if the assembly from the PDB has the same composition as a complex in the Complex Portal. If so, we assign the name of the Complex Portal entry to the PDB assembly. For example, the PDB entry 4f3l consists of two chains (UniProt entries Q9WTL8 and O08785) that can be mapped to the Complex Portal entry, CPX-3225. Therefore, we can name this heterodimer “CLOCK-Bmal1 transcription complex” using the name from the Complex Portal. We use the UniProt recommended name for homomeric protein assemblies in the PDB since they comprise repeated units of the same subunit. For homomeric protein assemblies that consist exclusively of unmapped components, we use either the entity names or the antibody names to name these assemblies. For example, the PDB entry 1ivi is named as dihydrolipoamide dehydrogenase while the PDB entry 12e8 is named as IG-heavy chain and IG-light kappa chain.

**Figure 6.**
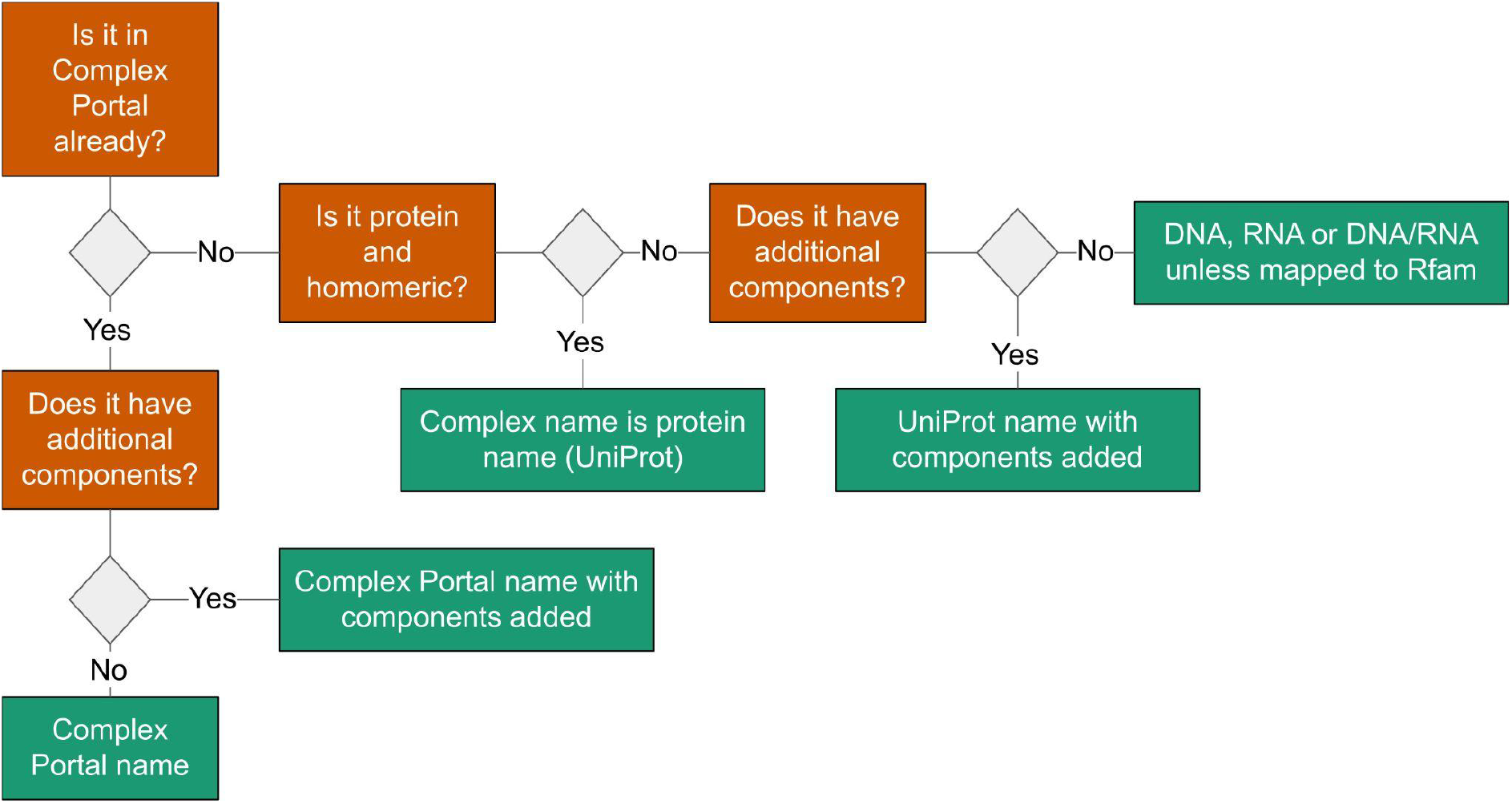
Decision tree for automated assembly descriptions. Our process uses a decision tree to automatically generate descriptions for assembly components based on data from external data resources and component categories.

However, many PDB assemblies have additional components not present in Complex Portal or UniProt entries. In such cases, we add names depending on the molecule type (i.e. DNA, RNA, or antibody). Nucleic acid assemblies that consist of a single component are named DNA, RNA, or DNA/RNA hybrid, depending on the polymer type of the assembly, unless they have components mapped to Rfam.

Most RNA-containing assemblies in the PDB are either ribosomes or spliceosomes, while the rest are mostly small assemblies that represent various small non-coding RNAs and tRNAs (52). Rfam is a valuable resource for identifying and naming RNA-containing assemblies in the PDB since it is a database of non-coding RNA families classified using multiple sequence alignments, consensus secondary structure, and covariance models (43). For example, there are 102 unique assemblies in the PDB consisting exclusively of components that can be mapped to Rfam. These include the S-adenosylmethionine (SAM) riboswitch (PDB 2gis), group-I intron (PDB 1grz), group-II intron (PDB 4y1n), and glucosamine-6-phosphate ribozyme (PDB 2gcv).

After the automated naming process, we check if the assembly is in the PDBe’s manually curated list of assembly names (Figure 7), which includes selected macromolecular complexes and their mapping to UniProt for each component. We also verify if all components in the assembly have a consistent Gene Ontology (GO) term associated with them (53, 54). For complex components, the GO term describes the whole complex, such as the nucleosome complex, where UniProt accessions for core histone proteins contain the GO term “nucleosome” (GO:0000786). If we can find a consistent GO cellular component term across all components in a complex, we use it to name the complex. If all components in the assembly have the same common GO cellular component term, we use that name as the assembly name. For instance, we named the assembly in PDB 2c35 “DNA-directed RNA Polymerase II” since it consists of two chains with the same common GO cellular component term.

**Figure 7.**
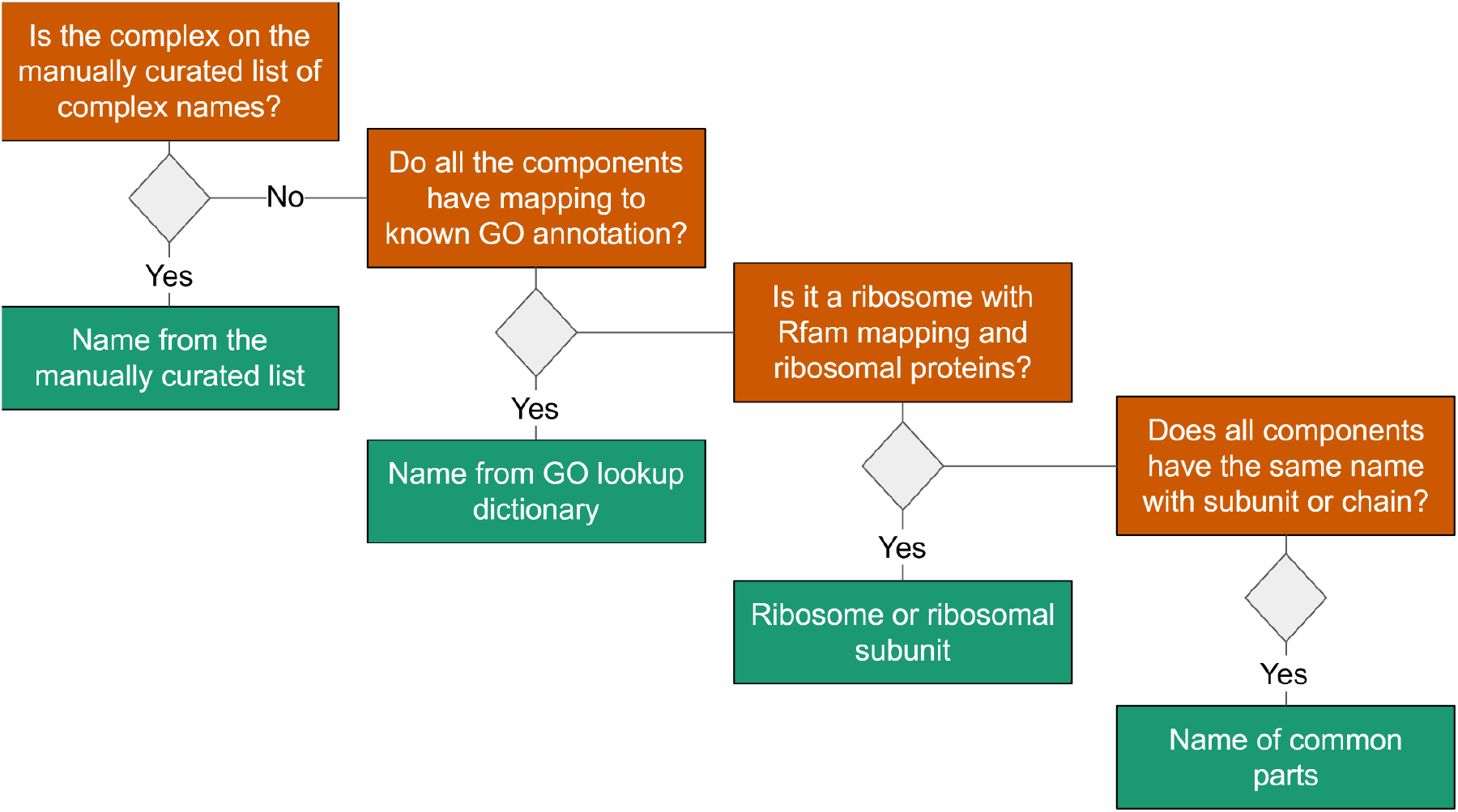
Decision tree for automated assembly naming. Our process attempts to assign human-readable complex names to all the unique composition descriptions. The process relies on a manually curated list of complex names, and when unavailable, it will look for data from GO annotations and common component names.

For ribosomal assemblies, we apply two additional steps. Since ribosomal composition can vary greatly between entries, we want to identify and name ribosomes even if they lack constituent protein or RNA components. First, we map rRNA components to Rfam to identify ribosomes. Then, we identify potential ribosomal assemblies by assessing if the assembly has both unmapped RNA molecules and ribosomal proteins.

## Data availability

The data pipeline described in this paper is integrated into the PDBe weekly release process to ensure that all new and updated PDB entries are assigned persistent complex identifiers. We provide public access to the data mapping between PDB and Complex Portal entries on our FTP area at https://ftp.ebi.ac.uk/pub/databases/pdbe-kb/complexes/, updated quarterly. Moreover, the code we used to analyse the data presented in this paper is available in a Jupyter notebook at https://github.com/PDBe-KB/pdbe-assemblies-analysis.

## Code availability

The source code for our data pipeline is publicly available on GitHub at https://github.com/PDBe-KB/process-complex-data. This repository contains a Python package that aggregates data for macromolecular complexes from the PDBe graph database, assigns unique identifiers, and generates human-readable names. Additionally, we have created a demo repository that allows developers to test and benchmark the complex identifier-generating process. This repository is available on GitHub at https://github.com/PDBe-KB/pdbe-complex-analysis-demo.

## Author contributions

S.D.A. and J.B. developed the complex identification process. S.D.A., S.N. and S.A. worked on integrating the process with the PDBe weekly release pipeline. S.N. developed the demo repository. S.D.A. and M.V. created the Jupyter notebook. R.G. created and maintained the manually curated list of complex names. M.V. and D.H. managed the project. I.P. designed the front-end changes, M.D. implemented them. M.V., J.E. and G.L.D. contributed to software development and testing. D.A. contributed to testing the open-source packages. S.G. contributed to the symmetry analysis. D.G. contributed annotations to Complex Portal. S.V. designed and developed the concept, secured funding and provided oversight. All authors contributed to the manuscript.

## References

1. Ramakrishnan, V. (2002) Ribosome Structure and the Mechanism of Translation. Cell, 108, 557–572.

2. Structure and mechanism of the RNA polymerase II transcription machinery.

3. Nooren, I.M.A. and Thornton, J.M. (2003) Diversity of protein–protein interactions. EMBO J., 22, 3486–3492.

4. Acuner Ozbabacan, S.E., Engin, H.B., Gursoy, A. and Keskin, O. (2011) Transient protein–protein interactions. Protein Eng. Des. Sel., 24, 635–648.

5. Raju, R.M., Goldberg, A.L. and Rubin, E.J. (2012) Bacterial proteolytic complexes as therapeutic targets. Nat. Rev. Drug Discov., 11, 777–789.

6. Hauser, A.S., Attwood, M.M., Rask-Andersen, M., Schiöth, H.B. and Gloriam, D.E. (2017) Trends in GPCR drug discovery: new agents, targets and indications. Nat. Rev. Drug Discov., 16, 829–842.

7. Lin, J., Zhou, D., Steitz, T.A., Polikanov, Y.S. and Gagnon, M.G. (2018) Ribosome-Targeting Antibiotics: Modes of Action, Mechanisms of Resistance, and Implications for Drug Design. Annu. Rev.Biochem., 87, 451–478.

8. Abrahams, J.P., Leslie, A.G.W., Lutter, R. and Walker, J.E. (1994) Structure at 2.8 Â resolution of F1-ATPase from bovine heart mitochondria. Nature, 370, 621–628.

9. Bowler, M.W., Montgomery, M.G., Leslie, A.G.W. and Walker, J.E. (2007) Ground state structure of F1-ATPase from bovine heart mitochondria at 1.9 A resolution. J. Biol. Chem., 282, 14238–14242.

10. Kabaleeswaran, V., Shen, H., Symersky, J., Walker, J.E., Leslie, A.G.W. and Mueller, D.M. (2009) Asymmetric Structure of the Yeast F1 ATPase in the Absence of Bound Nucleotides. J. Biol.Chem., 284, 10546–10551.

11. Xu, F., Wu, H., Katritch, V., Han, G.W., Jacobson, K.A., Gao, Z.-G., Cherezov, V. and Stevens, R.C. (2011) Structure of an agonist-bound human A2A adenosine receptor. Science, 332, 322–327.

12. Zhang, K., Zhang, J., Gao, Z.-G., Zhang, D., Zhu, L., Han, G.W., Moss, S.M., Paoletta, S., Kiselev, E., Lu, W., et al. (2014) Structure of the human P2Y12 receptor in complex with an antithrombotic drug. Nature, 509, 115–118.

13. Glukhova, A., Thal, D.M., Nguyen, A.T., Vecchio, E.A., Jörg, M., Scammells, P.J., May, L.T., Sexton, P.M. and Christopoulos, A. (2017) Structure of the Adenosine A1 Receptor Reveals the Basis for Subtype Selectivity. Cell, 168, 867–877.e13.

14. Groll, M., Ditzel, L., Löwe, J., Stock, D., Bochtler, M., Bartunik, H.D. and Huber, R. (1997) Structure of 20S proteasome from yeast at 2.4Å resolution. Nature, 386, 463–471.

15. Löwe, J., Stock, D., Jap, B., Zwickl, P., Baumeister, W. and Huber, R. (1995) Crystal structure of the 20S proteasome from the archaeon T. acidophilum at 3.4 A resolution. Science, 268, 533–539.

16. Schrader, J., Henneberg, F., Mata, R.A., Tittmann, K., Schneider, T.R., Stark, H., Bourenkov, G. and Chari, A. (2016) The inhibition mechanism of human 20S proteasomes enables next-generation inhibitor design. Science, 353, 594–598.

17. Ban, N., Nissen, P., Hansen, J., Moore, P.B. and Steitz, T.A. (2000) The Complete Atomic Structure of the Large Ribosomal Subunit at 2.4Å Resolution. Science, 289, 905–920.

18. Wimberly, B.T., Brodersen, D.E., Clemons, W.M., Morgan-Warren, R.J., Carter, A.P., Vonrhein, C., Hartsch, T. and Ramakrishnan, V. (2000) Structure of the 30S ribosomal subunit. Nature, 407, 327–339.

19. Yusupova, G., Jenner, L., Rees, B., Moras, D. and Yusupov, M. (2006) Structural basis for messenger RNA movement on the ribosome. Nature, 444, 391–394.

20. Karagöz, G.E., Duarte, A.M.S., Akoury, E., Ippel, H., Biernat, J., Morán Luengo, T., Radli, M., Didenko, T., Nordhues, B.A., Veprintsev, D.B., et al. (2014) Hsp90-Tau Complex Reveals Molecular Basis for Specificity in Chaperone Action. Cell, 156, 963–974.

21. Lapinaite, A., Simon, B., Skjaerven, L., Rakwalska-Bange, M., Gabel, F. and Carlomagno, T. (2013) The structure of the box C/D enzyme reveals regulation of RNA methylation. Nature, 502, 519–523.

22. Huang, C., Rossi, P., Saio, T. and Kalodimos, C.G. (2016) Structural basis for the antifolding activity of a molecular chaperone. Nature, 537, 202–206.

23. Rosenzweig, R., Moradi, S., Zarrine-Afsar, A., Glover, J.R. and Kay, L.E. (2013) Unraveling the Mechanism of Protein Disaggregation Through a ClpB-DnaK Interaction. Science, 339, 1080–1083.

24. Chua, E.Y.D., Mendez, J.H., Rapp, M., Ilca, S.L., Tan, Y.Z., Maruthi, K., Kuang, H., Zimanyi, C.M., Cheng, A., Eng, E.T., et al. (2022) Better, Faster, Cheaper: Recent Advances in Cryo–Electron Microscopy. Annu. Rev. Biochem., 91, 1–32.

25. Guaita, M., Watters, S.C. and Loerch, S. (2022) Recent advances and current trends in cryo-electron microscopy. Curr. Opin. Struct. Biol., 77, 102484.

26. Srivastava, A., Tiwari, S.P., Miyashita, O. and Tama, F. (2020) Integrative/Hybrid Modeling Approaches for Studying Biomolecules. J. Mol. Biol., 432, 2846–2860.

27. Kim, S.J., Fernandez-Martinez, J., Nudelman, I., Shi, Y., Zhang, W., Raveh, B., Herricks, T., Slaughter, B.D., Hogan, J.A., Upla, P., et al. (2018) Integrative structure and functional anatomy of a nuclear pore complex. Nature, 555, 475–482.

28. Chou, H.-T., Apelt, L., Farrell, D.P., White, S.R., Woodsmith, J., Svetlov, V., Goldstein, J.S., Nager, A.R., Li, Z., Muller, J., et al. (2019) The Molecular Architecture of Native BBSome Obtained by an Integrated Structural Approach. Structure, 27, 1384–1394.e4.

29. Aryal, R.P., Kwak, P.B., Tamayo, A.G., Gebert, M., Chiu, P.-L., Walz, T. and Weitz, C.J. (2017) Macromolecular Assemblies of the Mammalian Circadian Clock. Mol. Cell, 67, 770–782.e6.

30. wwPDB consortium (2019) Protein Data Bank: the single global archive for 3D macromolecular structure data. Nucleic Acids Res., 47, D520–D528.

31. Berman, H., Henrick, K., Nakamura, H. and Markley, J.L. (2007) The worldwide Protein Data Bank (wwPDB): ensuring a single, uniform archive of PDB data. Nucleic Acids Res., 35, D301–D303.

32. Lawson, C.L., Patwardhan, A., Baker, M.L., Hryc, C., Garcia, E.S., Hudson, B.P., Lagerstedt, I., Ludtke, S.J., Pintilie, G., Sala, R., et al. (2016) EMDataBank unified data resource for 3DEM. Nucleic Acids Res., 44, D396–D403.

33. Valentini, E., Kikhney, A.G., Previtali, G., Jeffries, C.M. and Svergun, D.I. (2015) SASBDB, a repository for biological small-angle scattering data. Nucleic Acids Res., 43, D357–D363.

34. Hoch, J.C., Baskaran, K., Burr, H., Chin, J., Eghbalnia, H.R., Fujiwara, T., Gryk, M.R., Iwata, T., Kojima, C., Kurisu, G., et al. (2023) Biological Magnetic Resonance Data Bank. Nucleic Acids Res., 51, D368–D376.

35. Burley, S.K., Kurisu, G., Markley, J.L., Nakamura, H., Velankar, S., Berman, H.M., Sali, A., Schwede, T. and Trewhella, J. (2017) PDB-Dev: a Prototype System for Depositing Integrative/Hybrid Structural Models. Struct. Lond. Engl. 1993, 25, 1317–1318.

36. Velankar, S., van Ginkel, G., Alhroub, Y., Battle, G.M., Berrisford, J.M., Conroy, M.J., Dana, J.M., Gore, S.P., Gutmanas, A., Haslam, P., et al. (2016) PDBe: improved accessibility of macromolecular structure data from PDB and EMDB. Nucleic Acids Res., 44, D385–D395.

37. Krissinel, E. and Henrick, K. (2007) Inference of macromolecular assemblies from crystalline state. J. Mol. Biol., 372, 774–797.

38. Ponstingl, H., Henrick, K. and Thornton, J.M. (2000) Discriminating between homodimeric and monomeric proteins in the crystalline state. Proteins Struct. Funct. Bioinforma., 41, 47–57.

39. Capitani, G., Duarte, J.M., Baskaran, K., Bliven, S. and Somody, J.C. (2016) Understanding the fabric of protein crystals: computational classification of biological interfaces and crystal contacts. Bioinformatics, 32, 481–489.

40. Duarte, J.M., Srebniak, A., Schärer, M.A. and Capitani, G. (2012) Protein interface classification by evolutionary analysis. BMC Bioinformatics, 13, 334.

41. Dey, S., Ritchie, D.W. and Levy, E.D. (2018) PDB-wide identification of biological assemblies from conserved quaternary structure geometry. Nat. Methods, 15, 67–72.

42. Dana, J.M., Gutmanas, A., Tyagi, N., Qi, G., O’Donovan, C., Martin, M. and Velankar, S. (2019) SIFTS: updated Structure Integration with Function, Taxonomy and Sequences resource allows 40-fold increase in coverage of structure-based annotations for proteins. Nucleic Acids Res., 47, D482–D489.

43. Kalvari, I., Argasinska, J., Quinones-Olvera, N., Nawrocki, E.P., Rivas, E., Eddy, S.R., Bateman, A., Finn, R.D. and Petrov, A.I. (2018) Rfam 13.0: shifting to a genome-centric resource for non-coding RNA families. Nucleic Acids Res., 46, D335–D342.

44. Meldal, B.H.M., Perfetto, L., Combe, C., Lubiana, T., Ferreira Cavalcante, J.V., Bye-A-Jee, H., Waagmeester, A., del-Toro, N., Shrivastava, A., Barrera, E., et al. (2022) Complex Portal 2022: new curation frontiers. Nucleic Acids Res., 50, D578–D586.

45. Ruan, Z., Orozco, I.J., Du, J. and Lü, W. (2020) Structures of human pannexin 1 reveal ion pathways and mechanism of gating. Nature, 584, 646–651.

46. Rodnina, M.V., Fischer, N., Maracci, C. and Stark, H. (2017) Ribosome dynamics during decoding. Philos. Trans. R. Soc. B Biol. Sci., 372, 20160182.

47. Zhou, J., Lancaster, L., Trakhanov, S. and Noller, H.F. (2012) Crystal structure of release factor RF3 trapped in the GTP state on a rotated conformation of the ribosome. RNA, 18, 230–240.

48. Pagès, G. and Grudinin, S. (2020) AnAnaS: Software for Analytical Analysis of Symmetries in Protein Structures. Methods Mol. Biol. Clifton NJ, 2165, 245–257.

49. Bhate, M.P., Molnar, K.S., Goulian, M. and DeGrado, W.F. (2015) Signal Transduction in Histidine Kinases: Insights from New Structures. Struct. Lond. Engl. 1993, 23, 981–994.

50. Wilkinson, M.D., Dumontier, M., Aalbersberg, Ij.J., Appleton, G., Axton, M., Baak, A., Blomberg, N., Boiten, J.-W., da Silva Santos, L.B., Bourne, P.E., et al. (2016) The FAIR Guiding Principles for scientific data management and stewardship. Sci. Data, 3, 160018.

51. Dunbar, J. and Deane, C.M. (2016) ANARCI: antigen receptor numbering and receptor classification. Bioinformatics, 32, 298–300.

52. Westhof, E. and Leontis, N.B. (2021) An RNA-centric historical narrative around the Protein Data Bank. J. Biol. Chem., 296.

53. Ashburner, M., Ball, C.A., Blake, J.A., Botstein, D., Butler, H., Cherry, J.M., Davis, A.P., Dolinski, K., Dwight, S.S., Eppig, J.T., et al. (2000) Gene Ontology: tool for the unification of biology. Nat.Genet., 25, 25–29.

54. The Gene Ontology Consortium (2021) The Gene Ontology resource: enriching a GOld mine. Nucleic Acids Res., 49, D325–D334.

